# Evolution of stable reciprocity through the maintenance of discrimination

**DOI:** 10.1101/2024.12.17.628959

**Authors:** Gilbert Roberts

## Abstract

Human cooperation among non-kin has long been accepted as being based on reciprocity – the idea that we return help from individuals that we re-meet. However, this paradigm has been challenged by Efferson et al. (Efferson, C., et al., Super-additive cooperation. *Nature*, 2024. 626, 1034-1041) who found that repeated interaction was insufficient to support cooperation without selection between groups. The authors based their conclusion on a model in which cooperative partners used a novel function to determine their response. In their system, cooperation through repeated interaction broke down. I point out here that the collapse of cooperation was not the result of the novel strategy set. Instead, it reflects a standard modelling result in which a cooperative environment allows drift towards unstable, undiscriminating strategies. I show how modifying model assumptions such as changing the implementation of mutation or allowing persistent defection maintain the variation which selects for discrimination. This discrimination in turn stabilizes reciprocity. I conclude that Efferson et al.’s results can readily be explained by existing theory and that repeated interaction can support cooperation without requiring between-group competition.

---

Reciprocity is a classical explanation for cooperation involving repeated interactions despite each move offering a temptation to defect (Trivers, 1971). It works when discriminating strategies, most famously ‘Tit-for-Tat’ (Axelrod & Hamilton, 1981), cooperate with cooperators but defect on defectors. However, this scenario has an Achilles heel. If cooperative strategies are profitable over multiple rounds, all agents will tend to evolve to cooperate through some form of reciprocation. In this cooperative utopia, the discrimination that makes strategies like Tit-for-Tat successful in an uncooperative environment becomes redundant and unconditionally cooperative strategies can drift in by mutation (‘indirect invasion’ (van Veelen, 2012)). This paves the way for uncooperative strategies to invade by exploiting the underlying naivety (Bendor & Swistak, 1995). Such dynamics “are the underlying basis for every model of cooperation” (Wahl & Nowak, 1999b).

Now, Efferson et al. (Efferson et al., 2024) have presented a model of repeated interactions in which cooperation collapses and have interpreted this as meaning that reciprocity cannot work on its own. They begin by describing a familiar scenario. In a cooperative population, individuals tend to start with high offers; this means there is little scope for escalation within a fixed endowment and consequently the parameters determining the degree of escalation in their model have little effect. Relieved from selection, the escalation parameters drift into a range where they are vulnerable to what they call ‘ambiguous’ strategies which engage in a form of ‘short-changing’ (Roberts & Sherratt, 1998). As a result, cooperation declines to low levels. However, the difference between Efferson et al.’s paper and previous work is that rather than offering the standard interpretation, the authors instead claim to have uncovered a flaw in the long-standing paradigm of repeated interaction as a pillar of cooperation. They further claim that uncovering this flaw was a consequence of modelling an ‘extended’ strategy space. Here, I show how the breakdown observed by Efferson et al. is due to cooperators not being exposed to the realistic levels of variability that are known to maintain the discriminating strategies that in turn provide stability.

## Methods

Simulation methods were based on those described in Efferson et al. and were coded independently in Microsoft Visual C. Simulations were initialized by setting up a population of 100 individuals. Play involved selecting every possible pair in turn (100*99/2 = 4950 pairings) and then for each pair, playing an iterated asynchronous continuous Prisoner’s Dilemma game for 100 rounds. On the first move, the leader was determined randomly. The leader donated part of its 1 unit of endowment according to its genetic strategy *x*, which was in increments of 0.01 between 0 and 1. The follower donated according to its response rule (*d-a*)*x*, where *a* and *d* were the left and right intercepts of a linear function of *x*. In subsequent rounds, both partners used this function to determine their response based on what their partner had donated. The difference between standard reciprocity and Efferson et al.’s modified version of reciprocity is explained in Figure 1.

**Figure 1.**
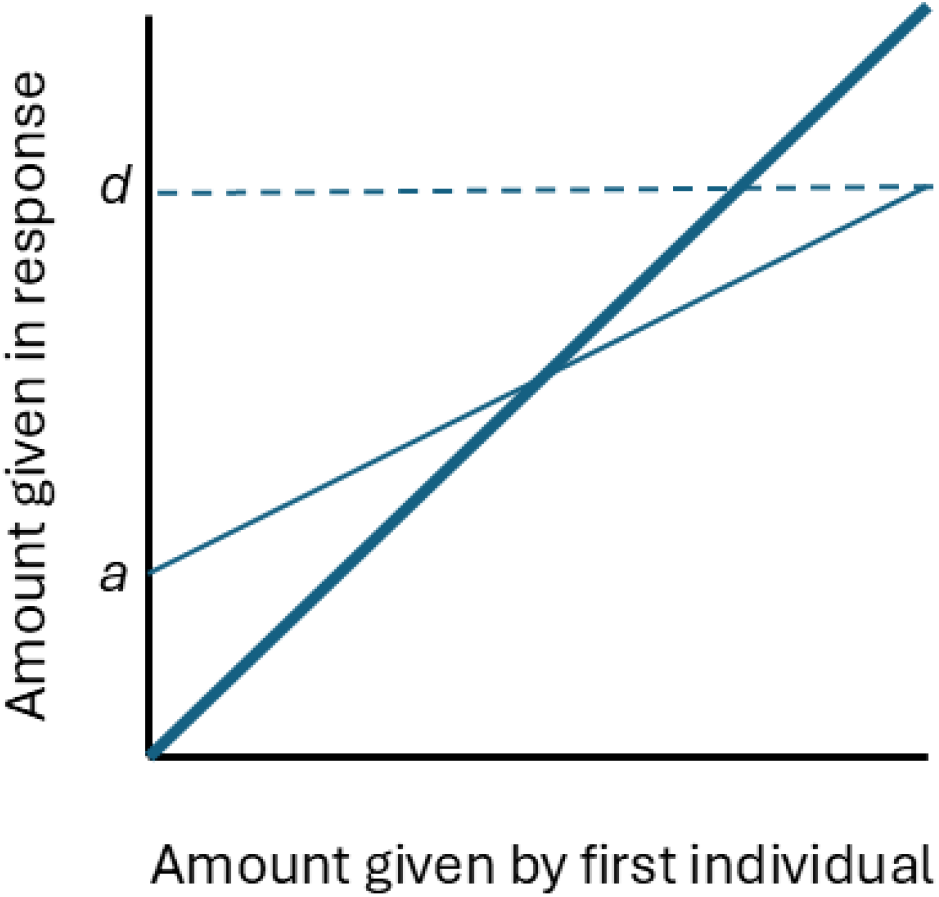
Reciprocal response functions. In classical reciprocity theory, the amount donated by any individual is a simple multiple of that received so can be represented by the bold line. In Efferson et al.’s expanded framework, a reciprocator donates according to its response rule (*d-a*) *x*, where *a* and *d* are the left and right intercepts of a linear function of the x axis amount donated by the first player and where investments are within the space 0,1.

**Figure 2.**
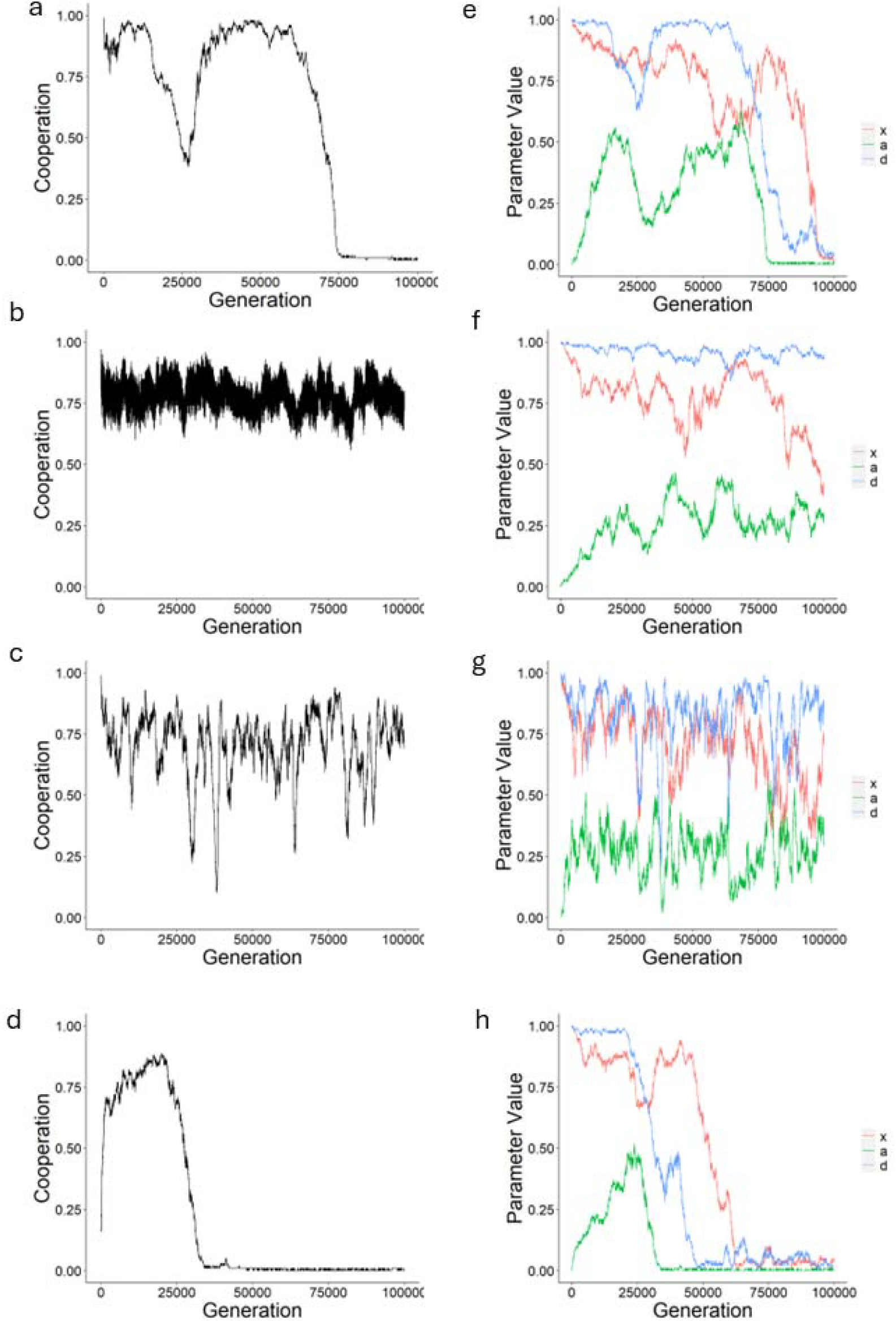
Evolutionary dynamics. In each case, the first of the ten simulations summarized in Figure 2 is plotted. In (a-d), cooperation is plotted as the proportion of the endowment donated to the partner in each generation, averaged across all N=100 individuals. In (e-f), the parameters in Efferson et al.’s model are plotted for each generation, again averaged across all N=100 individuals. (a & e) Replication of Efferson *et al*.’s model; (b & f) with a fixed proportion of 2% of individuals being phenotypic defectors; (c & g) with an additional mutation process giving a 1% chance that offspring will play ‘always defect’ (invest zero); (d & h) with implementation errors on 2% of moves.

**Figure 3.**
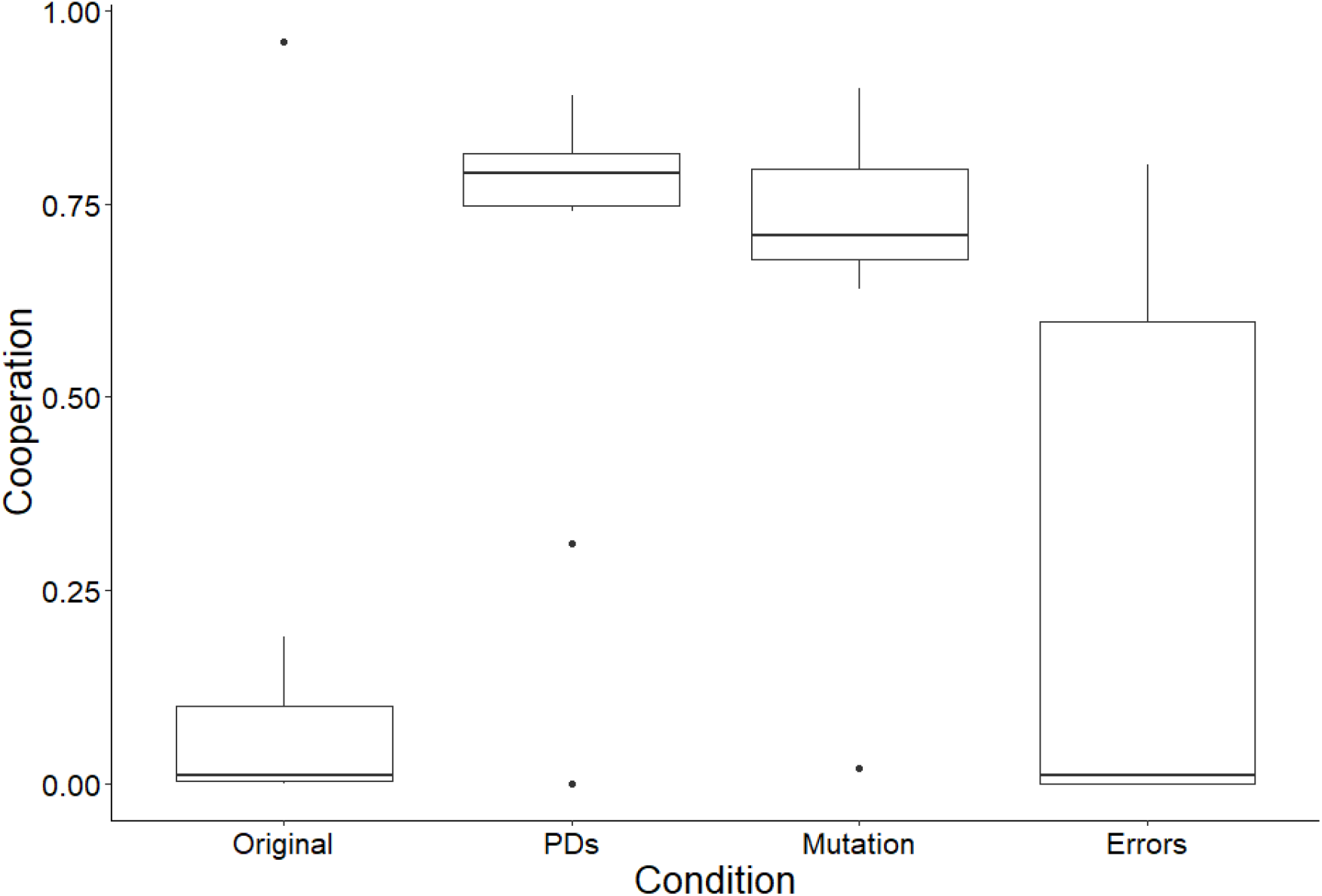
Boxplots summarizing cooperation levels across conditions. Plotted are the medians with inter-quartile ranges across 10 simulations for each of the conditions as in Figure 2, namely (1) as in the original Efferson et al. simulations; (2) with of 2% of individuals being phenotypic defectors; (3) with an additional mutation process giving a 1% chance that offspring will play ‘always defect’ (invest zero); and (4) with implementation errors on 2% of moves.

At the end of the full set of interactions, reproduction was proportional to payoffs and resulted in replacing the population of *N*=100 individuals. Following reproduction, mutation occurred. Mutation in Efferson et al. was implemented by adding or subtracting small increments to a strategy. In our model the genetic strategies increased or decreased in steps of 0.01 with probability 0.02 within the bounds of 0 to 1. However, this meant that in a cooperative environment, cooperators were not tested against defectors but just against those giving slightly less.

Simulations began with full cooperation given by *x*=1, *a*=0, *d*=1. Simulations ran for 100,000 generations of strategy evolution. I calculated cooperation as the average proportion of the 1 unit of endowment invested, averaged over the 4950 pairings * 100 rounds * 2 in a pair = 990,000 moves in each of 100,000 generations.

## Results

I began by coding a simple model using their response function to replicate the collapse of cooperation reported in Efferson et al. Note that the collapse of cooperation in Figure 1(a) can be understood in terms of the drift upwards of parameter *a* which presents an amount given to a partner independent of how much has been received (see Figure 1 (d)). This unconditional cooperation drifts so high that it destabilizes reciprocal cooperation, as seen in the collapse of the parameter *d* in Figure 1(d) and correspondingly in the rate of cooperation in Figure 1(a). This sequence warrants comparison with Figure 1 in (Sherratt & Roberts, 2001) which illustrates how the probability of cooperating with a defector can drift upwards until it comes to a point where it destabilizes cooperation. Reciprocity depends upon discrimination and when that is eroded it can no longer work.

A traditional way of overcoming evolution towards uniform cooperation in models has been to introduce persistent defection in the form of ‘phenotypic defectors’. These agents carry a cooperative genetic strategy yet their phenotype is defection (Lotem et al., 1999; Sherratt & Roberts, 2001). The rationale behind this is that there will always be some individuals who perhaps through a lack of resources are unable to cooperate. Phenotypic defectors were implemented in the model by replacing the genetic strategy with a phenotypic strategy of always defecting for 2% of individuals in each generation. Introducing these phenotypic defectors resulted in a high and stable level of cooperation (Figure 1(b)). Looking at the evolutionary dynamics of the response function parameters, it is clear that whilst there is some drift, we do not see the same increase in *a* that triggers the collapse of *d* in the case of Efferson et al.’s basic model. This result illustrates how allowing a small proportion of individuals that always defect has the paradoxical effect of increasing cooperation. Analytically, this has long been understood: discriminators do better against those who always defect because they suffer less exploitation than unconditional cooperators (Sherratt & Roberts, 2001). These discriminating strategies form the basis of stable cooperation, in contrast to the unstable cooperation of undiscriminating strategies.

A second way of ensuring there is the variation which feeds discrimination is to modify the assumptions about how mutation works. Efferson et al.’s model allows for continuously varying degrees of cooperation, but it is problematic in that their local mutation mechanism could mean that agents were tested only against those who gave a little more or less and not against full defectors. The test of cooperation is how well it stands up to defection. I therefore ran the model with an additional mutation process in which there was a 1% chance of each of the response parameters mutating to zero, thereby implementing an ‘always defect’ strategy. This resulted in a high level of stable cooperation (Figure 1(b)). Again, the evolutionary dynamics of the parameters show that there is not the destabilizing effect of a drift upwards of the *a* parameter (Figure 1(f)). Whilst one can discuss the merits of assuming local mutation versus mutation to giving nothing, the point is that the results presented by Efferson et al are fragile with respect to one of the key parameters of their model. Finally, to make clear that we are talking about variation in relation to discrimination and not errors of the type considered by e.g. (van Veelen, 2012), I examined the effect of introducing implementation errors by replacing 2% of responses with random responses. This treatment failed to stop the collapse of cooperation as observed in the original model (Figures 1(d) and (h)).

## Discussion

I find that the collapse of cooperation in Efferson et al.’s models is consistent with that of previous models of reciprocity and that it similarly disappears if selection for discrimination is maintained. Their results are therefore not the consequence, as they claim, of using a model with more response function parameters. Their claim that previous authors have found reciprocity only because they have excluded their strategies “by fiat” is unfair. Similarly, it cannot be concluded, as suggested by (Mathew, 2024) that their use of using a continuous model has uncovered previously unknown weaknesses of reciprocity. After all, “subtle cheating” (Trivers, 1971) or “short-changing” (Roberts & Sherratt, 1998) strategies are not new (see also (Wahl & Nowak, 1999a, 1999b)).

The significance of our analysis is in the fact that it overturns Efferson et al.’s claims about the evolution of human cooperation. Their thesis was that reciprocity did not work on its own but that cooperation could work if there was a positive interaction (in their terms a “super-additive” effect) with between-group competition. Their work therefore fit a narrative in which between-group competition has been a decisive process in human social evolution (Richerson et al., 2016). Others have gone further and interpreted the flawed models as demonstrating the role of cultural-group selection in human uniqueness (Mathew, 2024), the argument being that this process facilitates the spread of cooperation in humans relative to other animals. In fact, we cannot draw any such conclusions from Efferson et al.’s work. However, if reciprocity does work as I show, the evolution of human society may after all be explained by individuals cooperating because it is in their own long term best interests.

## Code and data availability

Code is available at osf.io/tw4gs

## Author contributions

G.R. conceived, modelled and wrote the manuscript.

## Ethics statement

This work did not involve any procedures involving humans or other animals.

## References

Axelrod, R., & Hamilton, W. D. (1981). The evolution of cooperation. Science, 211, 1390–1396.

Bendor, J., & Swistak, P. (1995). Types of evolutionary stability and the problem of cooperation. Proceedings of the National Academy of Sciences, 92(8), 3596–3600. 10.1073/pnas.92.8.3596

Efferson, C., Bernhard, H., Fischbacher, U., & Fehr, E. (2024). Super-additive cooperation. Nature, 626(8001), 1034–1041. 10.1038/s41586-024-07077-w

Lotem, A., Fishman, M. A., & Stone, L. (1999). Evolution of cooperation between individuals. Nature, 400(6741), 226–227.

Mathew, S. (2024). Why reciprocity is common in humans but rare in other animals. Nature, 626, 955–956. 10.1038/d41586-024-00308-0

Richerson, P., Baldini, R., Bell, A. V., Demps, K., Frost, K., Hillis, V.,… Newson, L. (2016). Cultural group selection plays an essential role in explaining human cooperation: A sketch of the evidence. Behavioral and Brain Sciences, 39, e30.

Roberts, G., & Sherratt, T. N. (1998). Development of cooperative relationships through increasing investment. Nature, 394(6689), 175–179.

Sherratt, T. N., & Roberts, G. (2001). The role of phenotypic defectors in stabilizing reciprocal altruism. Behavioral Ecology, 12, 313–317.

Trivers, R. L. (1971). The evolution of reciprocal altruism. Quarterly Review of Biology, 46, 35–57.

van Veelen, M. (2012). Robustness against indirect invasions. Games and Economic Behavior, 74(1), 382–393. 10.1016/j.geb.2011.05.010

Wahl, L. M., & Nowak, M. A. (1999a). The continuous prisoner’s dilemma: I. Linear reactive strategies. Journal of Theoretical Biology, 200(3), 307–321. 10.1006/jtbi.1999.0996

Wahl, L. M., & Nowak, M. A. (1999b). The Continuous Prisoner’s Dilemma: II. Linear Reactive Strategies with Noise. Journal of Theoretical Biology, 200(3), 323–338. 10.1006/jtbi.1999.0997

